# SATINN: An automated neural network-based classification of testicular sections allows for high-throughput histopathology of mouse mutants

**DOI:** 10.1101/2022.04.16.488549

**Authors:** Ran Yang, Alexandra Stendahl, Katinka A. Vigh-Conrad, Madison Held, Ana C. Lima, Donald F. Conrad

**Affiliations:** Division of Genetics, Oregon National Primate Research Center, Oregon Health and Science University, Portland OR 97006

## Abstract

**Motivation:** The mammalian testis is a complex organ with a hierarchical organization that changes smoothly and stereotypically over time in normal adults. While testis histology is already an invaluable tool for identifying and describing developmental differences in evolution and disease, methods for standardized, digital image analysis of testis are needed to expand the utility of this approach.

**Results:** We developed SATINN (Software for Analysis of Testis Images with Neural Networks), a multi-level framework for automated analysis of multiplexed immunofluorescence images from mouse testis. This approach uses a convolutional neural network (CNN) to classify nuclei from seminiferous tubules into 7 distinct cell types with an accuracy of 94.2%. These cell classifications are then used in a second-level tubule CNN, which places seminiferous tubules into one of 7 distinct tubule stages with 90.4% accuracy. We further describe numerous cell- and tubule-level statistics that can be derived from wildtype testis. Finally, we demonstrate how the classifiers and derived statistics can be used to rapidly and precisely describe pathology by applying our methods to image data from two mutant mouse lines. Our results demonstrate the feasibility and potential of using computer-assisted analysis for testis histology, an area poised to evolve rapidly on the back of emerging, spatially-resolved genomic and proteomic technologies.

**Availability and implementation:** Scripts to apply the methods described here are available from http://github.com/conradlab/SATINN.

## Background

Spermatogenesis is a cyclical process in mammalian seminiferous tubules that, under normal circumstances, results in continuous sperm production. Deficiencies in this complex but essential evolutionary process often result in male infertility, which is characterized by a reduction or complete absence of mature sperm count (Schlegel 2004) and currently affects approximately 1 in 7 couples worldwide (Agarwal, et al. 2021). While key regulatory genes (Krausz and Casamonti 2017) (Mou, et al. 2013), chromosomal microdeletions (Tiepolo and Zuffardi 1976) (Ma, et al. 1992), and environmental factors (Gabrielsen and Tanrikut 2016) have been associated with cases of male infertility, another 30% to 40% cases remain idiopathic (Nieschlag, Behre and S 2010), indicating that our current knowledge of the molecular machinery of spermatogenesis (Chen, et al. 2018) (Wang, et al. 2018) (Guo, et al. 2017) (Shami, et al. 2020) (Suzuki, Diaz and Hermann 2019) is far from complete.

Histology is the premier method for phenotyping spermatogenic defects and has set the foundation of understanding spermatogenesis in several key organisms, including humans (Clermont 1966) (Paniagua and Nistal 1984), non-human primates (Clermont and Lebland 1959) (Clermont and Antar 1973), and mice (Yoshida 2008) (Yoshida 2012) (Nakata 2019). A single cross-section of a mouse testis contains approximately 120 tubules that house tens of thousands of germ cells discriminated by histological markers into 20 cell types (Chiarini-Garcia and Meistrich 2008) (Drumond, Meistrich and Chiarini-Garcia 2011) and 12 tubule stages (Oakberg 1956) (Russell, et al. 1990) (Ahmed and de Rooij 2009) that serve as landmarks in the cycle of spermatogenesis. But while the quality and quantity of testis histology has greatly improved over the last decade, computational tools capable of handling and analyzing that data are just emerging. Traditional histology is low-throughput due to the time-consuming nature and expertise required to manually analyze the images. Clinical histopathology typically focuses on identifying only a handful of severe phenotypes, such as Sertoli cell only, germ cell maturation arrest, and hypospermatogenesis (Abdullah and Bondagji 2011) (Hentrich, et al. 2011), which only offer insights at a coarse resolution.

To enable higher level analyses and to increase data processing efficiency, we aim to integrate histology with both computational image processing and machine learning. The potential of computational processing of histological images has been shown by 3D modeling of seminiferous tubules in rat, mouse (Nakata, Wakayama and Sonomura, et al. 2015) (Nakata, Sonomura and Iseki 2017) (Nakata 2019), and Syrian hamster (Nakata, Yoshiike, et al. 2021), to better understand the physical constraints of these systems. Machine learning itself has shown useful applications in other fields such as cancer research, where gene expression-based neural networks can distinguish between several cancer cell types (Mostavi and Chiu 2020) and image recognition networks can detect breast cancer cells based on changes in actin filament structure (Oei, et al. 2019). However, adapting these learning algorithms to analyze testis histology remains largely unexplored, apart from a single recent study from (Xu, et al. 2021) which used a neural network to stage Hematoxylin and Eosin (H&E)-stained tubule cross-sections.

Our goal in this study is to develop and assess a computational method to evaluate histopathology using automated classification of mouse seminiferous cell types and tubule stages from immunofluorescence (IF) images. To our knowledge, this report is the first of a publicly available, neural network-based classification method for IF testis images, which have unique features for the computer to learn from. Our workflow has the benefit over similar methods of making no assumptions about the composition of cells within tubules, which reduces processing times and enables functionality under non-ideal conditions, such as for meiotic arrest mutants which lack entire cell type populations. It also opens the door to extensive network refinement by using fluorescent markers with additional specificity, as well as downstream quantification of those markers of interest, something that would be more difficult to do using traditional immunohistochemistry (IHC) stains.

In this paper, we describe and validate our neural network trained to automatically classify mouse seminiferous cell types and tubule stages from IF images stained with a basic set of markers. We show that we are able to computationally recapitulate the previously described meiotic-arrest phenotype of *Mlh3*^*-/-*^ mice. Additionally, we use the high sensitivity of our software to make biological inferences on an undisclosed mouse mutant line that exhibits a much milder phenotype that would typically be impossible to detect and quantify by eye. We conclude by discussing the implications of our work to understanding the mechanical limitations of spermatogenesis as well as the research and clinical potential of combining image recognition software with the field of infertility.

## Results

### Neural networks classify mouse seminiferous tubules and nuclei with above 90% accuracy

To facilitate high-throughput statistical analysis of seminiferous tubules with various genetic backgrounds, we developed SATINN (Software for Analysis of Testis Images with Neural Networks) to automate cell type and tubule stage classification (overview in Fig. 1).

**Fig. 1:**
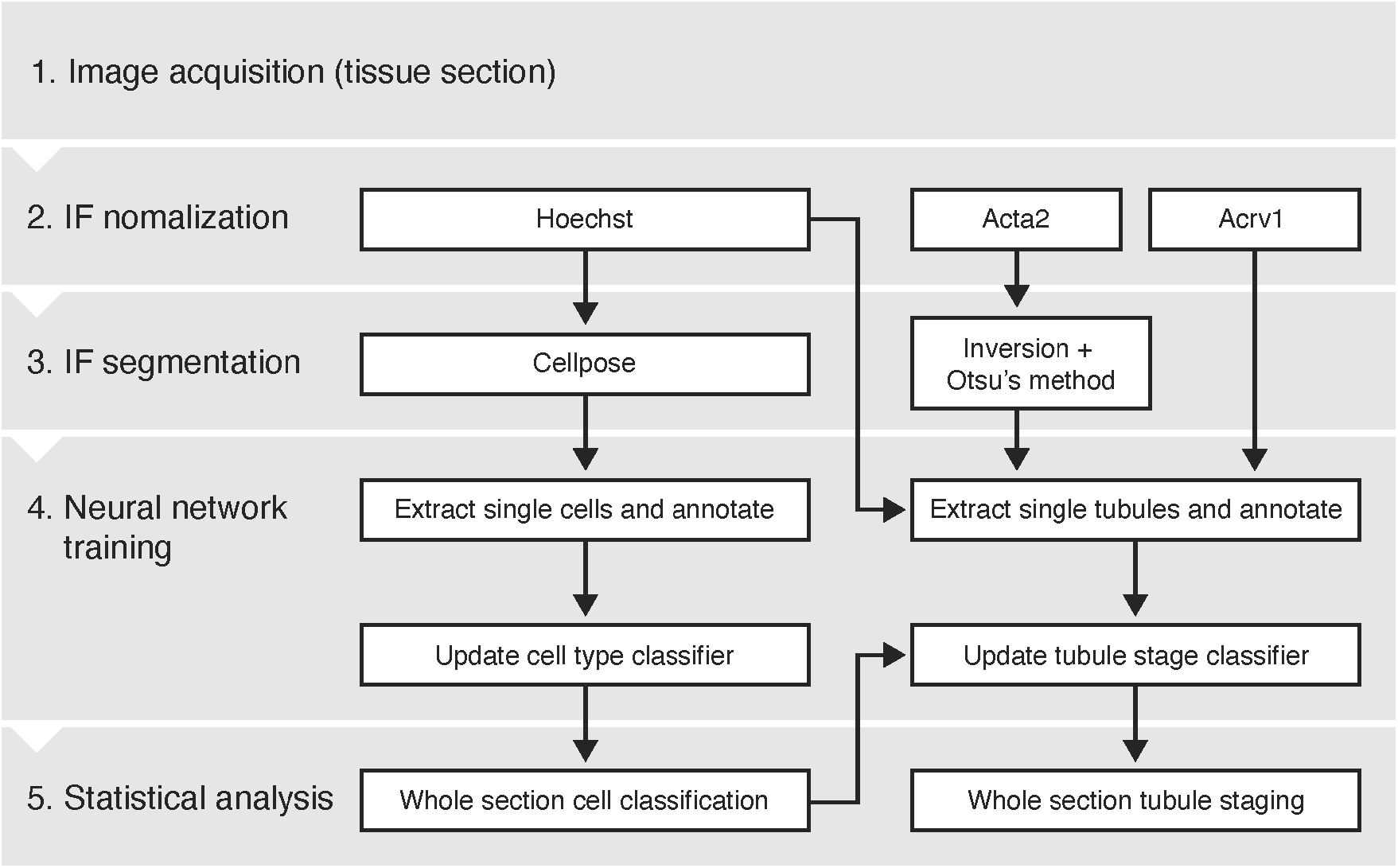
Overview of image processing. Raw images are acquired (see methods) and processed autonomously through this workflow, enabling high-throughput analysis of large numbers of nuclei per image (each whole testis cross-section from mouse contains approximately 120 tubules, or the order of hundreds of thousands of nuclei).

We acquired cross-sectional images of mouse seminiferous tubules (see methods and supplemental methods) containing the following color channels: Hoechst (a nuclear marker), Actin Alpha 2 (Acta2), and Acrosomal vesicle protein 1 (Acrv1). Acta2 and Acrv1 were used to assist the tubule classifier as described below. We segmented cell nuclei using Cellpose (Stringer, et al. 2021) and tubules using Otsu’s method (Otsu 1979), automated object extraction, and manually annotated over 7,800 cells and 2,000 tubules to use for neural network training. We then built two convolutional neural network (CNN) classifiers using MATLAB®’s Deep Learning Toolbox: a cell type classifier based on seven different cell types (overview in Fig. S1) and a tubule classifier with seven tubule stages. Finally, we evaluated the accuracy of our neural networks against their own training data and manually annotated validation data.

To train our cell type classifier, we used 7,871 representative images of nuclei from seven different cell types: round (rSPD), intermediate (iSPD), and elongated (eSPD) spermatids; primary (SPC) and secondary (SPCII) spermatocytes; spermatogonia (SPG); and Sertoli cells annotated from 88 wildtype seminiferous tubules (11 testis sections from 6 animals). We found that using only the Hoechst channel for training was sufficient to achieve a classification accuracy of 84.1% across all cell types (Fig. 2A).

**Fig. 2:**
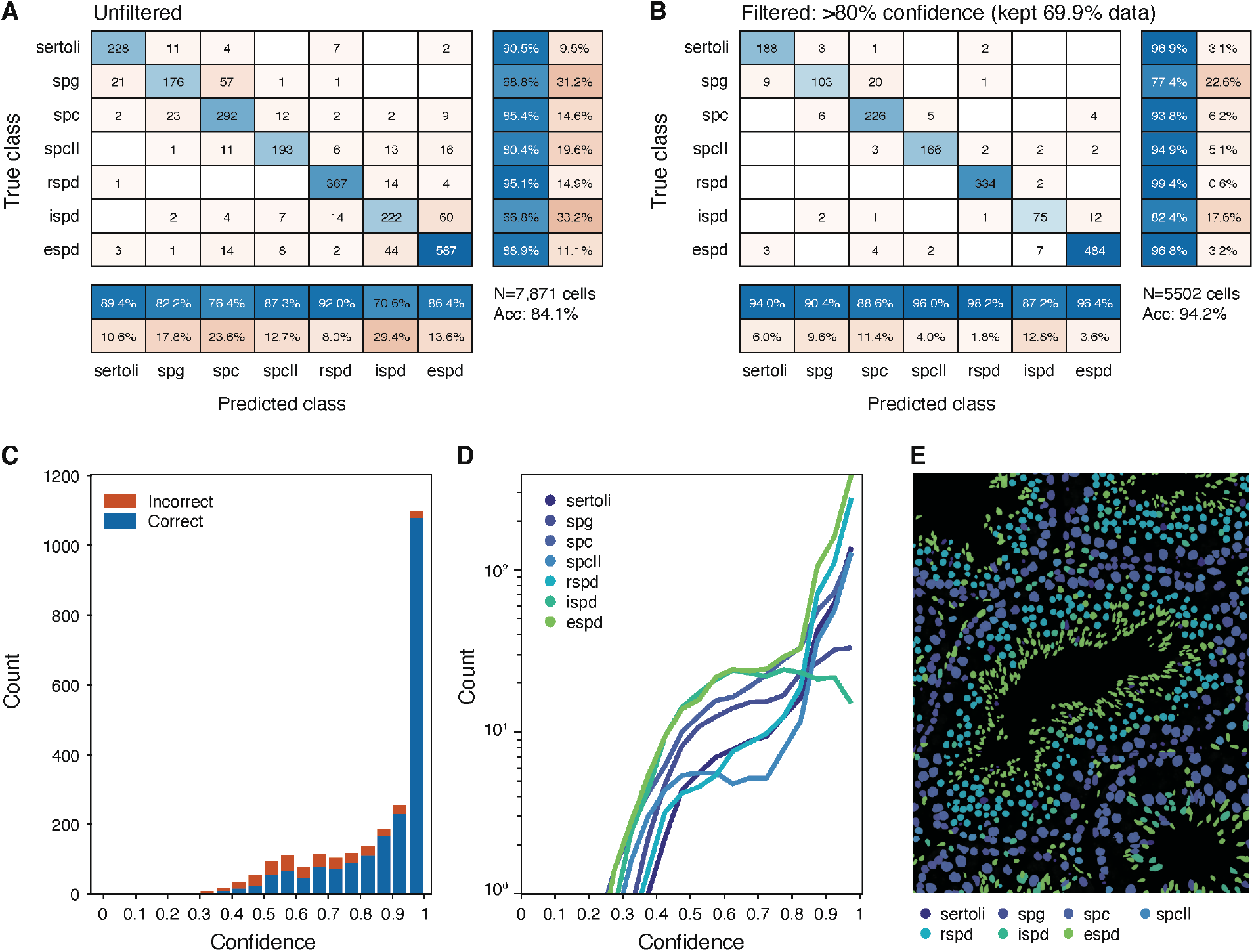
Results of cell type classification (training data validation). We used 7,871 annotated images to train the neural network in figure 1B to recognize seven different cell types. 70% of this data was randomly selected as the training set while the remaining 30%, presented here, was used to determine the NN efficacy. (A) Confusion matrix. Rows indicate annotated cell classes; columns indicate NN predictions. Bottom two rows of percentages indicate positive predictive values and false discovery rates respectively, while right two columns of percentages indicate true positive and false positive rates respectively. Overall accuracy of the test data was 85.6%. (B) High confidence confusion matrix. The same matrix as in (A), but predictions with confidence less than 80% were not included, retaining 72.7% of the data. The overall accuracy increases to 94.2%. (C) Histogram of confidence with all cells pooled into bins corresponding to their assignment confidence at 5% intervals. The highest confidence intervals also contain the highest total counts and the highest fraction of correct calls within those intervals. (D) Classifier confidence per cell type. All cell types have a high count of high confidence calls. However, some developmentally adjacent cell types such as spermatogonia and spermatocytes have some low confidence calls, as expected due to attempting to discretize a continuous biological process. (E) Visual close-up of a tubule with cell type predictions. Each colored object represents a segmented cell; color indicates the cell type. N: sample size; Acc: Accuracy

Filtering out low confidence calls (LCF, defined as below 80% confidence) resulted in 94.2% of training images being classified correctly while retaining a majority (69.9%) of the data (Fig. 2B). We confirmed the validity of thresholding in this way using two metrics: first, we observed that lower confidence calls have the highest fractions of incorrect classifications and vice versa (Fig. 2C); and second, each cell type has similar confidence distributions (Fig. 2D). Classification of nuclei from a whole wildtype section was visibly accurate (Fig. 2E). Though we could have increased classification accuracy by including additional cell type-specific information such as Acrv1 to demarcate eSPDs, we prioritized simplicity of imaging requirements by minimizing the number of necessary markers and applicability of our neural network to non-wildtype tubules that may not express supplementary markers in the same way or at all.

Similar to the cell type classifier, our tubule stage classifier was trained using a set of 2,037 representative images (24 testis sections from 15 mice) corresponding to various tubule stages. These training data required Acta2 to isolate individual tubules and Acrv1 to more readily distinguish developmentally adjacent stages. Histologists often use a 12-stage classification system to describe the arrangement of cells within the tubules (Ahmed and de Rooij 2009). To improve the performance of our approach, we grouped similar stages together, resulting in a seven-stage classifier. We incorporated cell class predictions to further assist the tubule classifier, input as seven additional image layers of probability for each cell detected within a tubule. This resulted in a general classification accuracy of 80.0% (Fig. 3A), which increased to 90.4% (Fig. 3B) post-LCF. As with the cell classifier, we ensured that the majority of calls were high confidence (Fig. 3C) and that no tubule stages in particular were more difficult to classify than any other (Fig. 3D). The result of tubule stage classification on an unannotated wildtype section is shown in figure 3E.

**Fig. 3:**
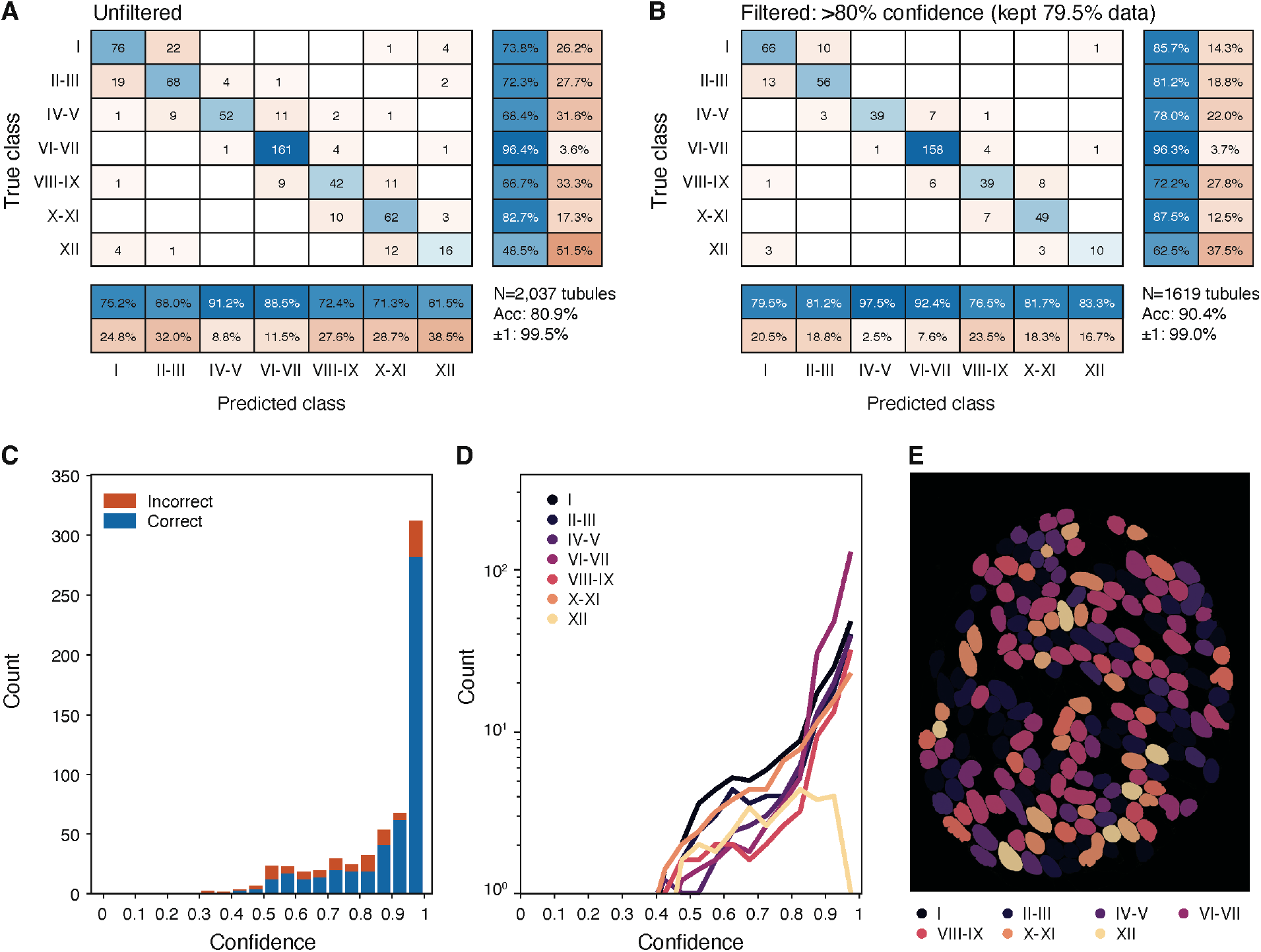
Results of tubule stage classification (training data validation). We used 2,037 annotated tubules to train a neural network similar to that shown in figure 1B to recognize seven different tubule stages. The methods and plots are similar to those described in figure 2. (A) Confusion matrix. The overall accuracy of tubule staging is 80%, increased to 96.7% given one degree of error (e.g. stage I can be classed as stage II-III or stage XII). (B) High confidence confusion matrix. The overall accuracy increases to 90.4% and nearly all calls (99.0%) are within one degree of error. (C) Histogram of confidence at 5% intervals with all tubule stages pooled. (D) Classifier confidence per tubule class. The stage XII class has notably fewer high confidence calls, likely due to being easily confused with stage I. (E) Visual overview of tubule segmentation and stage classification. Each object is colored by its predicted stage class. N: sample size; Acc: Accuracy. ±1: Accuracy within one adjacent class error.

### Properties of wildtype nuclei are reproducibly measured and are resolved at the level of individual stages

In addition to using training data to validate our neural networks, we used unannotated wildtype tubules (representative brightfield image in Fig. 4A) to further confirm the functionality and accuracy of our classification. We first compared our CNN-derived nuclei quantification with cell counts from published literature. Despite some variability between references (e.g. (Tegelenbosch and de Rooij 1993) (Oakberg 1956) (Yang and Oatley 2014) (Clermont 1972)), our counts were overall consistent with established values, and we show one such comparison here with (Nakata, Wakayama and Takai, et al. 2015). Our total segmented nuclei count per tubule was 297 ± 123 (values hereafter reported as mean ± st.dev) sourced from over 2,400 tubules (left panel on Fig. 4B). While this number is about 35% lower than the reference report due to differences in methodology (we use a nuclear marker and computerized segmentation as opposed to manual cell counting in an H&E image), the relative proportions of each cell population remain comparable (right panel on Fig. 4B).

**Fig. 4:**
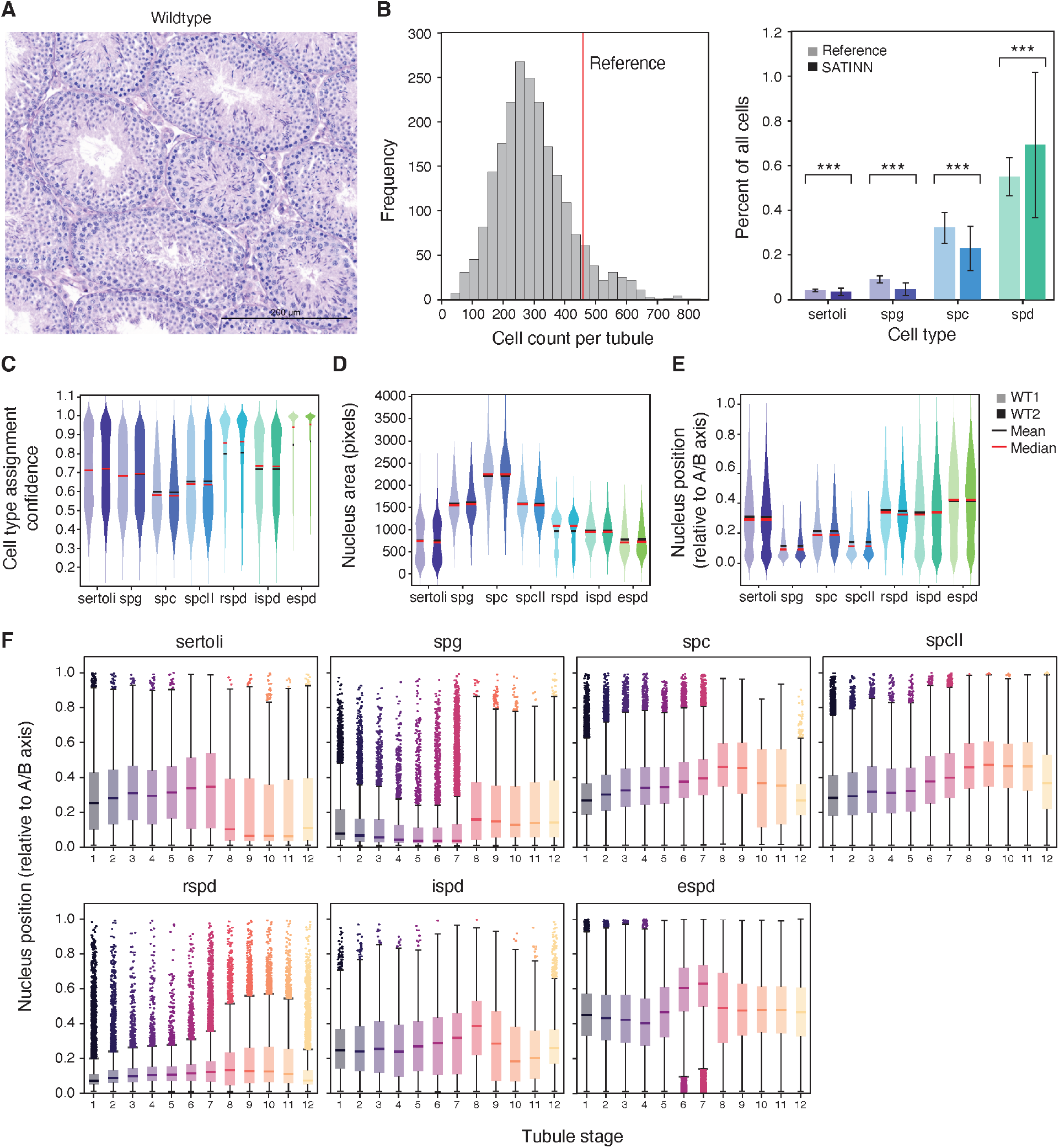
Statistical analysis of an independent wildtype dataset (not used for training or validation). (A) Representative brightfield image of a wildtype tubule (shown: *Mlh3+/+*). (B) Cell counts detected by our workflow, compared to (Nakata, Wakayama and Takai, et al. 2015). (left) Total cell counts per tubule; (right) Normalized ratios of specific cell types. (C-F) Cross-sections of whole testes (unannotated) from five mice were imaged and split into two subgroups. Each subgroup contains an image from each mouse. Within each cell type pair (two adjacent violin plots), the left distribution originates from subgroup 1; right from subgroup 2. Each distribution was processed using our custom quantile normalization method (Fig. S2). Each pair is not significant (p > 0.01) unless noted. (C) Cell type assignment confidence, arranged by cell type. (D) Nuclear area in pixels. (E) Apical-basal position (ABP). This custom metric is defined as the location of a nucleus’ centroid relative to its tubule centroid and the segmented tubule edge. (F) ABP sorted by tubule stage and cell type.

Batch effects are a common concern with high-throughput experiments. We found subtle but significant differences among biological replicates of wildtype samples in classification confidence scores, presumably due to technical variables impossible to account for (reagent batch, operator technique, etc.). To address these batch effects, we developed a modified version of quantile normalization (Fig. S2) that could be used with our data, and found this approach removed all differences in nuclei features between two wildtype subgroups (Mann-Whitney U-test p > 0.01; Fig. 4C-E).

Having validated the accuracy of our cell types and tubule stages, we developed a number of quantitative measures that summarize the properties of cell nuclei within the tubules, which, to our knowledge, have not been rigorously quantified, including area (Fig. 4D), apical-basal position (ABP, Fig. 4E) and relative orientation, i.e. the deviation of the object’s orientation from the apical-basal axis (Fig. S3A-D). These measurements reflect our current knowledge from testis biology. For instance, meiotic SPC and SPCII show the largest nuclei and, together with and pre-meiotic germ cells (SPG), were concentrated to the basal side of the tubule, while post-meiotic cells (spermatids) occupied apical positions. Normalized nuclear orientations did not have large magnitudes of change, though they slightly shifted towards a radial distribution (pointing toward lumen) as the cells progressed from SPCs to eSPDs (Fig. S3E).

To better visualize spatial rearrangements in seminiferous tubules that may not be immediately apparent at first glance, we plotted ABP (Fig. 4F) and relative orientation (Fig. S3F) with respect to tubule stage. We were able to recapitulate eSPDs moving into the lumen (more apical position) at stages VI-VII, while having an otherwise stable ABP distribution. On the other hand, rSPDs steadily increase in ABP as spermatogenesis progresses, reflecting their apical migration as they mature. Finally, we note that Sertoli ABP drops sharply (more basal) at stage VIII, after blood-testis barrier remodeling, while at the same time, classified SPG are shifted apically as they become committed to differentiation. Through this method, we validated many previously established observations of spermatogenesis in an unbiased way, which increased our confidence in our statistical methods and overall approach.

### Cell type clusters are identified by nearest neighbor mapping, and spermatogenic index fluctuates with tubule stage

To demonstrate higher level analytical capabilities of SATINN, we designed several statistical features which may be useful to studying seminiferous tubule biology. We began by quantifying spatial relationships among different cell types: for each classified cell type (reference), we found the nearest neighbors of every other cell type (targets) within that tubule (example shown in Fig. 5A). We counted and normalized the cell types that correspond to the single nearest target for each reference (Fig. 5B) and found strong correlations among spermatids (rSPD, iSPD and eSPD), as well as the Sertoli-SPG-SPC block, an accurate reflection of the tissue’s architecture. We additionally analyzed radial distances between nearest neighbors, which ignores the circumferential component in order to establish apical-basal directionality between two nuclei (Fig. 5C). Consistent with our understanding of the spatial organization of these cell types, most spermatid targets averaged negative (basal) values while SPG targets averaged positive (apical). Finally, we evaluated the efficacy of spermatogenesis by calculating the spermatogenic index of each tubule, which we defined as the ratio of eSPD to SPG count. Assuming ideal meiotic conditions this value should be at least 4 (Hess and de Franca 2008). When we plotted spermatogenic index in wildtype tubules with respect to stage (Fig. 5D), we found this value fluctuates depending on the tubule stage but is indeed centered around 4, dropping at stage VIII, when eSPDs are released into the lumen through spermiation.

**Fig. 5:**
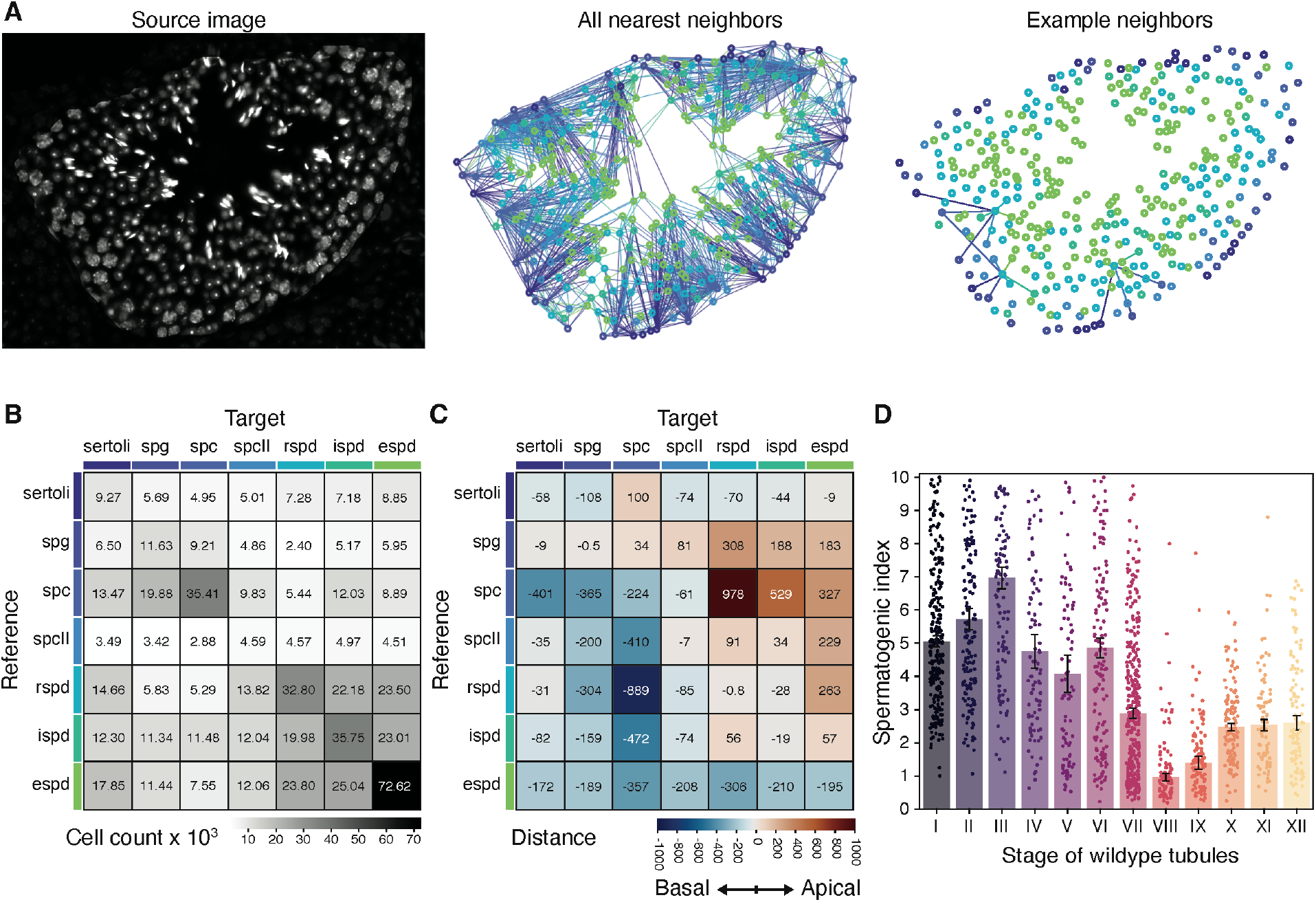
Higher order statistical analysis of an independent wildtype dataset. (A) Nearest neighbors analysis. (left) Sample source image of cells in a single tubule. Cells outside of the tubule boundary are dimmed. (center) Full network of nearest neighbors detected in source image. Circles represent cell centroids; their colors represent their cell type. Lines are drawn from each cell (reference) to the nearest neighbor of every cell type (targets) and are color-coded based on the target’s type. (right) Sample network from three arbitrarily chosen cells, shown for clarity. (B) Mean target-reference pair counts per tubule, from single nearest neighbor networks. Values shown in grid are x103. (C) Mean target-reference distances. (D) Spermatogenic index (ratio of eSPD:SPG) of wildtype tubules sorted by annotated stage.

### Application of our workflow in histopathology quantification of a spermatogenesis-deficient mutant, Mlh3

To test the applicability of SATINN for histopathology analysis, we used a well-characterized mutant strain with severe defects in germ cell development. Mlh3 is a DNA mismatch repair protein that has been shown to be essential for meiotic recombination during spermatogenesis (Lipkin, et al. 2002) (Toledo, et al. 2019). Homozygous *Mlh3*-deficient (*Mlh3*^*-/-*^) mice are sterile, as SPCs are unable to complete meiosis, resulting in tubules with maturation arrest and a lack of late and post-meiotic germ cells (Fig. 6A). From three *Mlh3*^*-/-*^ cross-sections (350 tubules and 41,000 nuclei), we find that the total nuclei count in each tubule was reduced by 74.8% compared to wildtype (data not shown), which is consistent with the overall testis size reduction observed by (Lipkin, et al. 2002) and others. The primary contributing factor to the nuclei count reduction was the loss of classified spermatids (55.2% reduction of eSPD count; 89.5% of rSPD, data not shown), in agreement with classification from single-cell RNA-seq data (Jung, et al. 2019). To address the cell types in *Mlh3*^*-/-*^ that were classified with an improbable cell type (i.e. spermatids and SPC-II), we filtered those classes and re-normalized the remaining distributions (i.e. of Sertoli cells, SPC, and SPG, Fig. S4A). A comparison among data management methods (removal, adjustment, and no filtering) of improbable cell classes did not appreciably change the results (not shown). Our statistical analysis found little to no variation in nuclear size and orientation, as expected (Fig. 6B and S4B, respectively). However, our computational method revealed a reversal in the spatial organization of Sertoli cells from SPGs and SPCs (Fig. 6C), as well as increasing cell density along the basal tubule edge (Fig. S4C).

**Fig. 6:**
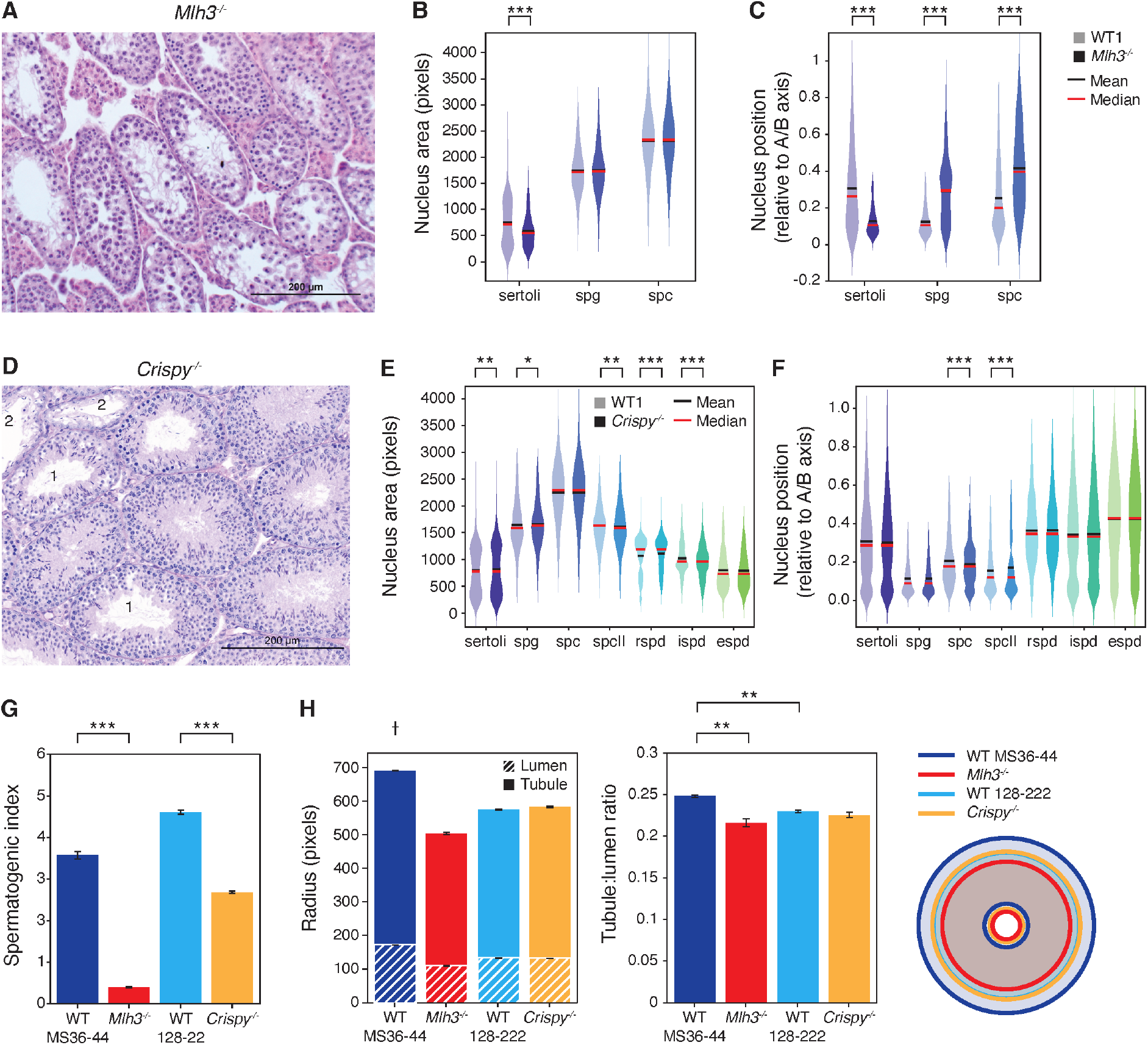
Statistical analysis of mutant cells. (A-C) *Mlh3-/-*. (A) Representative brightfield image of *Mlh3-/-* mutant seminiferous tubules. Three whole testis sections were derived from two mice and pooled into one subgroup (right plot in each colored pair) and plotted against the “WT1” subgroup (from Fig. 4, left plot in each colored pair). Each pair is not significant (p > 0.01) unless noted. Mlh3 cells that were not classified as Sertoli, spermatocyte, or spermatogonia were omitted for this analysis. (B) Cell area in pixels, split by cell type. (C) ABP. (D-F) *Crispy-/-*. (D) Representative brightfield image of *Crispy-/-* mutant seminiferous tubules. Six whole testis sections were derived from three *Crispy-/-* mice and pooled into one subgroup and plotted against the “WT1” subgroup, similar to that of Fig. 5. Each colored pair is not significant (p > 0.01) unless noted. (E-F) Metrics plotted are of those found in (B-C, respectively). (G) Spermatogenic index (ratio of eSPD:SPG) sorted by genotype. *Mlh3-/-* values are provided pre-filtering, as the spermatogenic index would otherwise be zero by definition. (H) Measurements of tubule and lumen radius. (left) Bar plots indicating tubule radius (higher value) and lumen radius (lower value). (center) The calculated tubule-to-lumen ratio. (right) Visual representation of mean circularized tubules for each genotype.

### Detection of subtle morphological changes in Crispy mutants

The hardest challenge in histopathology is the identification of minor changes undetectable by the human eye. Therefore, we tested SATINN on a mutant with a milder phenotype to calibrate its ability to detect subtle differences. To do this, we analyzed *Crispy*^*-/-*^ mutants (Fig. 6D), a mouse line with a targeted 5kb deletion of an evolutionarily conserved non-coding sequence that is predicted to impact spermatogenic gene expression (Okhovat *et al*., manuscript in preparation). Analyzing six *Crispy*^*-/-*^ cross-sections (1,164 tubules and 320,000 cells) we found subtle but significant changes in the nuclear areas of most cell types (Fig. 6E) and an apical shift in nuclei location of SPC and SPCII (Fig. 6F, p < 10^−5^) without disruption in other cell types.

Lastly, we performed tubule-level analysis. The spermatogenic index (Fig. 6G, ratio of eSPD:SPG counts) averaged close to 4 for both our annotated wildtype data and *Crispy*^*+/+*^ tubule populations (labeled as WT MS36-44 and WT 128-222 respectively in Figs. 6G-H). Despite the majority of *Crispy*^*-/-*^ tubules appearing morphologically indistinguishable to wildtype, they had a notably lower spermatogenic index, averaging slightly less than 3 across all tubules (p < 10^−5^). This finding, which would not have been apparent in a qualitative evaluation, may provide useful insight on the functional mechanisms of this mutant. We also calculated tubule and lumen radii (Fig. 6H) for the genotypes used in this paper. Wildtype tubules were larger than the *Crispy*^*+/+*^, which is likely a result of differences in fixation protocols for those equivalent genotypes. Nonetheless, *Mlh3*^*-/-*^ tubules remained the smallest (p < 10^−5^), in agreement with known literature (Toledo, et al. 2019). Additionally, when comparing the following datasets which were fixed with the same protocol, *Crispy*^*-/-*^ mutants are the same size as *Crispy*^*+/+*^ (p > 0.64), suggesting that unlike *Mlh3*, the *Crispy* mutation does not affect tubule size. A brief calculation of the ratio between lumen and tubule radii revealed that these proportions remain roughly similar across all tubules, with the largest differences once again being attributed to the wildtype dataset, though the significance values are barely below threshold (p = 0.006).

## Discussion

In this paper we present SATINN, a software that performs a high-throughput analysis of immunofluorescence images from whole mouse testis cross-sections. We apply the image recognition capabilities of convolutional neural networks to the field of reproductive biology, resulting in automated detection and classification of thousands of nuclei into 7 cell types and hundreds of tubules into 7 stages of spermatogenesis, from a single cross-section image with very high accuracy. We show the benefit of collecting large amounts of data for the otherwise inaccessible exploration of fine spatial relationships between tissue structures, and how they can contribute to a better understanding of testicular biology. Importantly, we demonstrate that this software can be used to recapitulate known histopathologies of the *Mlh3*^*-/-*^ mouse mutants and detect mild phenotypic alterations in the histology of an unpublished mouse mutant, *Crispy*^*-/-*^. To our knowledge, this code is the first of its kind to be publicly available and will be immediately useful to analyze IF images of mouse testis generated with the same staining schema used here (Hoechst, Acta2, Acrv1).

Recently (Xu, et al. 2021) began to explore the potential of automated classification of tubule stages. However, this work was limited to brightfield images, which inherently lacks the capability of simultaneously detecting multiple molecular targets, and did not expand on the identification of tubular cell-types, which are crucial for the study of complex biological processes in the testis. Our approach has improved on these limitations through the classification and statistical analysis of more precise cell types and tubule stages, which we then applied to studying mutant morphologies. While we demonstrated SATINN’s ability to detect and quantify subtle morphological changes, it was necessary to make careful interpretations due to the limitations of image recognition software. As with all machine learning methods that use discrete classes to categorize a continuous biological process like spermatogenesis, we expect the presence of cell type or tubule stage intermediates to arise as errors during classification. To mitigate this effect, we used a post-classification filtering based on classifier confidence (LCF), which improved the accuracy of both classifiers, and importantly, did not compromise our statistical power due to the large volume of cells and tubules that can be analyzed from each image. Additionally, due to the highly sensitive nature of CNNs, we expected the presence of batch effects and addressed them by quantile normalization (Fig. S2). We also noticed that specific measurements could be affected by experimental conditions. Different fixatives vary in the degree of distortion produced in the tissues during fixation (Mortensen and Brown 2003), which is reflected in our data by the significantly larger tubule radius of wildtype tubules fixed with 4% PFA (WT MS36-44) when compared to those fixed with modified Davidson’s fixative (WT 128-222; Fig. 6H). These differences could have important implications when assessing fertility of mutant mice, emphasizing the need to compare wildtype and mutant samples processed with the same experimental conditions.

Our current challenges include the difficulty of extrapolating what the neural network has learned to unknown states of pathology, including other mutant phenotypes. As we have already optimized what our classifiers can learn from our current datasets (analytics not shown), the next step would be to diversify training sources or to annotate state-specific images from other mutations, through a more streamlined training process if necessary. Further improvements of our current workflow to more accurately model spermatogenesis could include replacing the discrete system with a continuous or cyclical one. Additionally, tubule segmentation currently requires data from the Acta2 marker. This could potentially be overcome by using spatial cues, such as peritubular cell identification, to delimit the tubule boundaries. Integration of other molecular markers of interest would be useful in refining existing classification and enabling quantification of relevant developmental phenomena. Continued adaptation of these computational methods will ensure more reproducible and efficient image processing, along with identification of higher order features. Finally, the use of this software requires some degree of proficiency in MATLAB®, which is proprietary software, and will be addressed by the development of a graphic-user interface to improve accessibility.

Although the work presented here was done in mouse testes, we developed the workflow with as few assumptions as possible to enable further potential applications. A major benefit of this work is to assist fertility research in model animals, including mice, humans, and non-human primates. Adaptation of SATINN in other organisms such as humans will undoubtedly present additional challenges, such as reconciliation of the multiple tubule stages present in a single human seminiferous tubule cross-section. However, the rewards for overcoming these challenges would be greatly impactful, as the number of stages per cross-section is correlated with efficiency of spermatogenesis (Chaturvedi and Johnson 1993) and could therefore be used as a proxy of fertility assessment. Furthermore, the ability to detect multiple proteins or mRNAs of interest is one of the hallmarks of fluorescence immunoassays. With this software, we provide the means for high-throughput analysis of molecules *in situ*, with the spatial context that only histological images can confer. One potential avenue to pursue would be to integrate automated image analysis with emerging spatial omics technologies and help bring unprecedented refinement in our ability to assess complex molecular pathways. In the long term, these improved computerized image analysis methods hold the promise of automating the analysis of testicular biopsies, a task which currently requires laborious manual evaluations performed by extensively trained experts. Prospective clinical applications of automated testis image analysis range from to the diagnosis and prognosis of assisted human reproductive technologies (McLachlan, et al. 2006) (Faes, et al. 2013) (Esteves, Roque and Garrido 2018), treatment and prevention of testicular diseases (Elsherbeny and Abdelhay 2019), and even risk management of fertility in COVID-19 patients (Yang, et al. 2020).

## Methods

### Experimental setup

To capture subtle image differences resulting from experimental variables, the samples used to train the neural network were processed under different conditions: a) fixation method – perfusion or immersion; b) fixative type – 4% Paraformaldehyde (PFA) or modified Davidson’s fixative and c) assay – Immunofluorescence with or without Tyramide signal amplification (TSA) and RNA fluorescence *in situ* hybridization followed by immunofluorescence (FISH-IF). The boundaries of the seminiferous tubules were detected with anti-ACTA2 (1:100, Santa Cruz, sc-32251), the staging of tubules was performed with anti-ACRV1 (1:200, Proteintech, 14040-1-AP) and Sertoli cell nuclei marked using anti-SOX9 (1:100, Sigma-Aldrich, HPA001758). See the supplemental Methods for a detailed description of the experiments.

### Imaging

Whole testis sections were scanned with an Olympus VS120 slide scanner using a 40X (NA 0.95; 0.17μm/pixel) objective and a BrightLine® Sedat filter set (Semrock, DA/FI/TR/Cy5/Cy7-5×5M-B-000). Adjacent sections on the same slide without primary antibody or probe were used as negative controls to set the threshold of laser intensity during acquisition.

### Image processing and segmentation

Images were processed autonomously using the following methods in MATLAB® unless otherwise noted. (1) Intensity normalization was done using a basic top-hat filter with disk structuring element of radius 100 (value empirically determined to be suitable for these datasets). (2a) Cell segmentation was done using the Hoechst channel in Cellpose (Stringer, et al. 2021) with expected size of 30 pixels. (2b) Tubule segmentation was done using the Acta2 channel. A small amount of dilation was used to join imperfect tubule outlines, followed by whole image opening in order to remove small amounts of interstitial space (non-tubule regions). Thresholding was determined by Otsu’s method (Otsu 1979) and the dilation factors were corrected post-segmentation.

### Convolutional neural networks and training validation

Convolutional neural networks (CNN) to classify cell type and tubule stage were designed in MATLAB® with similar architectures (Fig. S1). The input for cells was a 50×50 pixels normalized Hoechst image centered on the centroid of each segmented cell, whereas for tubules, a 2000×2000 pixels normalized image containing Hoechst, Acta2, and Acrv1 was used instead. Objects whose CNN input image exceeded the boundaries of the source image were padded with zeros. For cell nucleus CNN training, each image was manually annotated with its cell type. We chose to annotate intratubular cells only. The categorization of “intermediate spermatid” refers to the developmental state between round and elongated spermatids found around stage IX.

Individual cell images were sourced from 12 different testis sections to minimize bias and to acquire sufficient quantities of low-abundance cells, such as Sertoli. Likewise, for tubule CNN training, images were manually annotated with their tubule stages, though they were subsequently grouped into the categories seen in Fig. 3A, due to similarities between adjacent stages. Tubules were sourced from the same 12 sections used to source cells, and additional care was taken to acquire a sufficient number of low-abundance stages. 70% of each class of annotated data were randomly selected for CNN training purposes, with the remaining 30% used to validate the efficacy of each CNN. Multiple attempts to train CNNs resulted in validation accuracies within 2% variance. The custom-built code for creating these figures and analyzing the data is available at the GitHub repository (https://github.com/conradlab/SATINN).

### Statistical distributions and tables

Tables of summary statistics for nuclei and tubules were built with the following components: image sources, neural net outputs, and segmented object statistics. CNN outputs include probabilities of each class, while object statistics include the output of the MATLAB® function ‘regionprops’ as well as the following outputs of custom-built code: (1) normalized apical-basal position of nuclei was inferred from the distance between (a) the nucleus centroid and tubule centroid and (b) the nucleus centroid and nearest tubule edge; (2) relative nucleus orientation was determined by the minimum angle formed between the major axis vector of the nucleus and the vector formed between its centroid and the tubule centroid. A value close to the minimum (0) indicates orientation along the apical-basal axis (radial) whereas one close to maximum (90) indicates a circumferential (orthogonal) orientation.

### Quantile Normalization

We used a modified version of quantile normalization that was originally described by (Hicks and Irizarry 2015) in order to mitigate the impact of batch effects. Rather than quantile-normalizing a 2D matrix, we extended the design to a 3D matrix containing the following components: features (x-variable) are cell types; samples (y-variable) are the image source; and for each feature and image source, 10,000 randomly chosen observations are recorded in the z-direction. Each z-vector containing observations from a single feature-sample is sorted in ascending order, then each xy-frame is independently quantile-normalized using the same method as in (Hicks and Irizarry 2015). p-values are extracted by conducting either paired t-tests or Mann-Whitney U-tests for each feature between two samples. (See Fig. S2 for visual representation.)

## Acknowledgments

The authors would like to thank the Chang Lab at the Knight Cancer Institute, Oregon Health & Science University, especially Dr. Young Hwan Chang and Dr. Erik Burlingame for the valuable discussions and suggestions. The authors also thank the Carbone Lab at OHSU and the Ahituv laboratory at the University of California, San Francisco, for the animal samples provided, and members of the Conrad lab for helpful discussion.

This work was supported by funding from the National Institutes of Health (R01HD078641 and P50HD096723). Research reported in this publication was supported by the Office of the Director of the National Institutes of Health under award number P51OD011092 to the Oregon National Primate Research Center. The ONPRC Integrated Pathology Core provided support services for the research. The content is solely the responsibility of the authors and does not necessarily represent the official views of the National Institutes of Health.

