## Supplementary Material for "SATINN: An automated neural network-based classification of testicular sections allows for high-throughput histopathology of mouse mutants"

### Supplementary Information for Yang et al.

#### Supplementary Methods

*Tissue collection and staining*

All animal experiments were performed in compliance with the regulations of the respective host institutions. Tissue was collected from sexually mature mice (> 8 weeks old). *Mlh3*$^{-/-}$ mice (The Jackson Laboratory; #018845) were bred at the Washington University in Saint Louis and processed as previously described (Jung, et al. 2019). Mice of the Crispy line were bred by the Ahituv laboratory at the University of California, San Francisco, and fixed testes were received in 70% Ethanol. Wildtype C57BL/6J (The Jackson Laboratory; #000664) were bred in-house at the Small Laboratory Animal Unit (ONPRC/ OHSU). Mice were sacrificed by $CO_2$ asphyxiation, testes were collected and tunica albuginea was pierced with a 31 G needle. For a subset of the samples, mice were sedated with inhaled 3% isoflurane and transcardially flushed with 0.9% saline before being perfused with 4% Paraformaldehyde [PFA; Electron Microscopy Sciences (EMS), #50-980-487] in 1X Phosphate Buffered Saline (PBS, ThermoScientific, J75889-k8). Testes were fixed overnight at 4°C in either 4% PFA or modified Davidson's fixative (EMS, #64133-50). The next day, the tissue was either directly transferred into a 70% ethanol solution or washed in 1X PBS and dehydrated through a series of increasing ethanol concentrations up to 70% in preparation for paraffin embedding. Sections of 5 µm were used for all of the assays.

Testis were stained with hematoxylin and counter-stained with either Periodic acid-Schiff reagent (*Crispy*$^{-/-}$ testis; Thermo Scientific, #87007) following manufacturer's recommendations, or eosin (*Mlh3*$^{-/-}$), as previously described (Jung, et al. 2019).

*Fluorescence in situ hybridization with Immunofluorescence – FISH-IF*

FISH was performed using RNAscope® Multiplex Fluorescent Reagent Kit v2 [Advanced Cell Diagnostics (ACD), 323100] according to manufacturer's guidelines with 15 minutes of both protease plus treatment and target antigen retrieval. Instead of counterstaining with DAPI, slides were rinsed in deionized water (di$H_2O$) for immunofluorescence. Briefly, slides were incubated in blocking solution [10% normal donkey serum (Sigma-Aldrich, D9663), 1% BSA (Sigma-Aldrich, A9647-50G), 1X PBS] for 15 minutes, in primary antibody for 1 hour, in secondary antibody (1:500, Invitrogen, A21202 and A21207) for 20 minutes and in 1µg/mL Hoechst 33342 (Invitrogen, H3570) for 5 minutes. These incubations were performed at 37°C in a humid chamber in the HybEZ™ oven (HybEZ™ II Hybridization System; ACD, 321711).

*Immunofluorescence without TSA*

Tissue was deparaffinized in Xylenes and rehydrated through a series of decreasing concentrations of Ethanol. Tissue was permeabilized in 0.25% Triton X-100 (in 1X PBS), heat-induced antigen retrieval was performed in Universal Antigen Retrieval Reagent (Abcam, ab208572) and incubated in blocking solution (10% normal donkey serum, 1% BSA, 1X PBS). Primary antibodies were incubated over-night at 4°C, secondary antibodies (1:500, Invitrogen, A21202 and A21207) for 1 hour at room temperature and tissue was counterstained with 1µg/mL Hoechst 33342.

*Immunofluorescence with TSA*

Tissue was processed following the same steps as for FISH-IF but without protease treatment. Then, tissue was blocked in 1X PBS supplemented with 10% normal donkey serum and 1% BSA for 20 minutes and incubated in primary antibody for 1 hour. Two rounds of tyramide signal amplification were then performed using the Opal 4-color anti-rabbit manual IHC kit (Akoya Biosciences, NEL840001KT) with Opal 520 and Opal 570 fluorophores and anti-mouse HRP (1:500) and anti-rabbit HRP (1:500) secondary antibodies. Between rounds, RNAscope® Multiplex FL v2 HRP blocker was used to stop the amplification. There was no need to strip primary antibodies since they were raised in different species. Antibody incubations were performed for 15-20 minutes at 37°C in a humid chamber in the HybEZ™ oven and TSA for 10 minutes at room temperature.

### Supplementary Figures

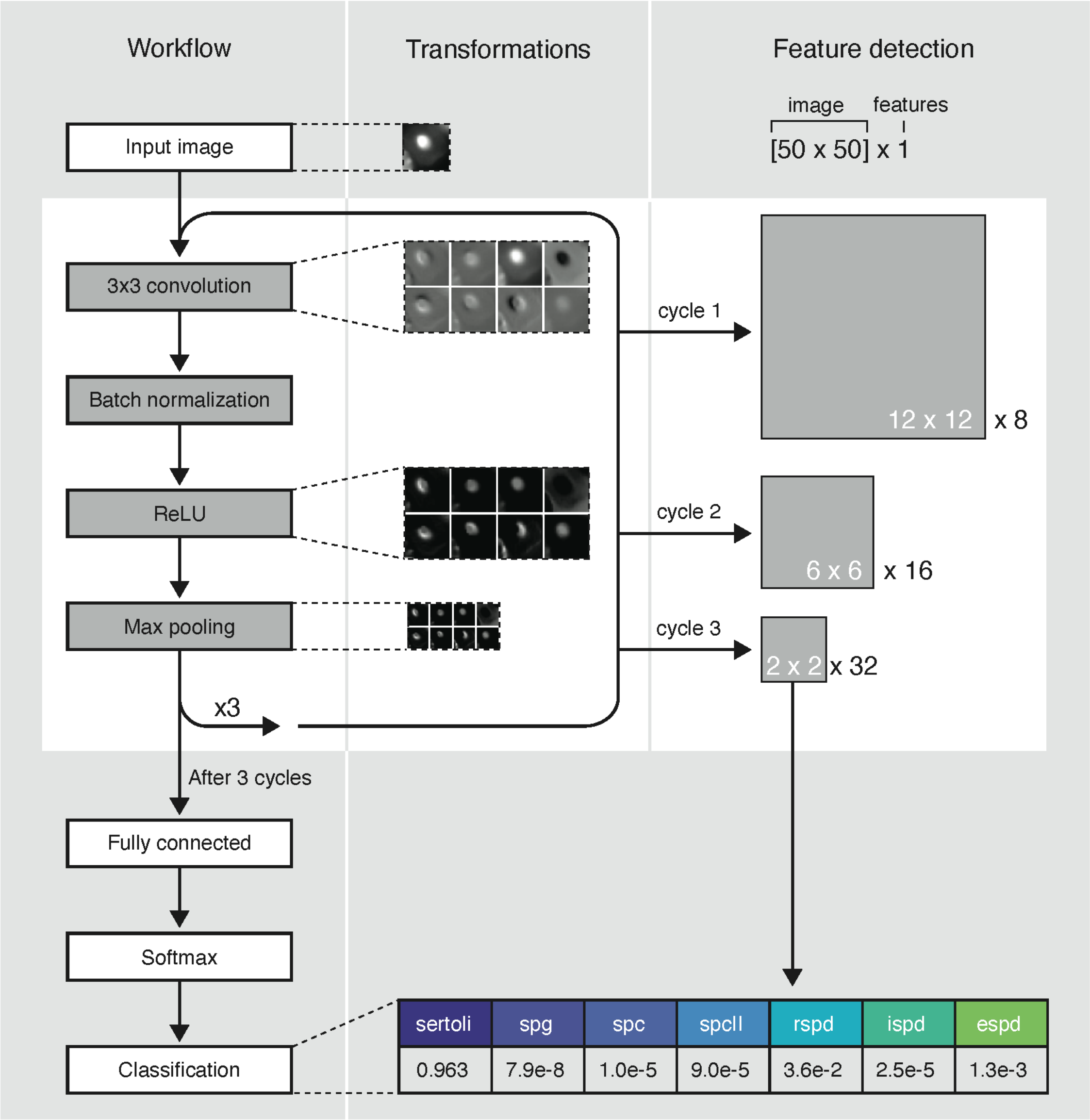


Fig. S1: Neural network workflow (cell classifier shown). A simple convolutional network was sufficient to achieve high accuracy classification of 7 different spermatogenic cell types. A sample image of a Sertoli cell is used to show the transformations leading to feature detection and predicted class. Overview of the convolutional neural network (CNN) architecture (cell classification). A 50x50 input image for each cell is used. Images show the intermediates of a

Sertoli cell after the first convolution, normalization, and pooling steps, resulting in an 8-feature map. Each additional cycle builds upon features found in the previous one, before outputting the probabilities of classification for each image. The NN identifies this Sertoli cell with 96.3% confidence, as indicated in the bottom table that shows classification probabilities for each cell class. Our NN for tubule classification is similar to what is shown here, with some modified parameters.

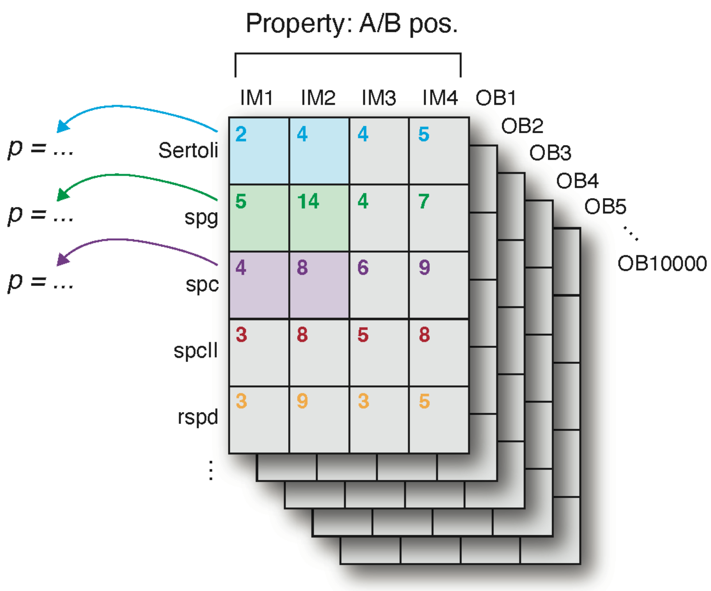


Fig. S2: Visual representation of statistical analysis using quantile-normalized data. After normalization (see methods), p-values are derived from statistical tests using observation vectors for each feature from pairs of images (each pair of colored squares represents two z-vectors from which the p-value is derived).

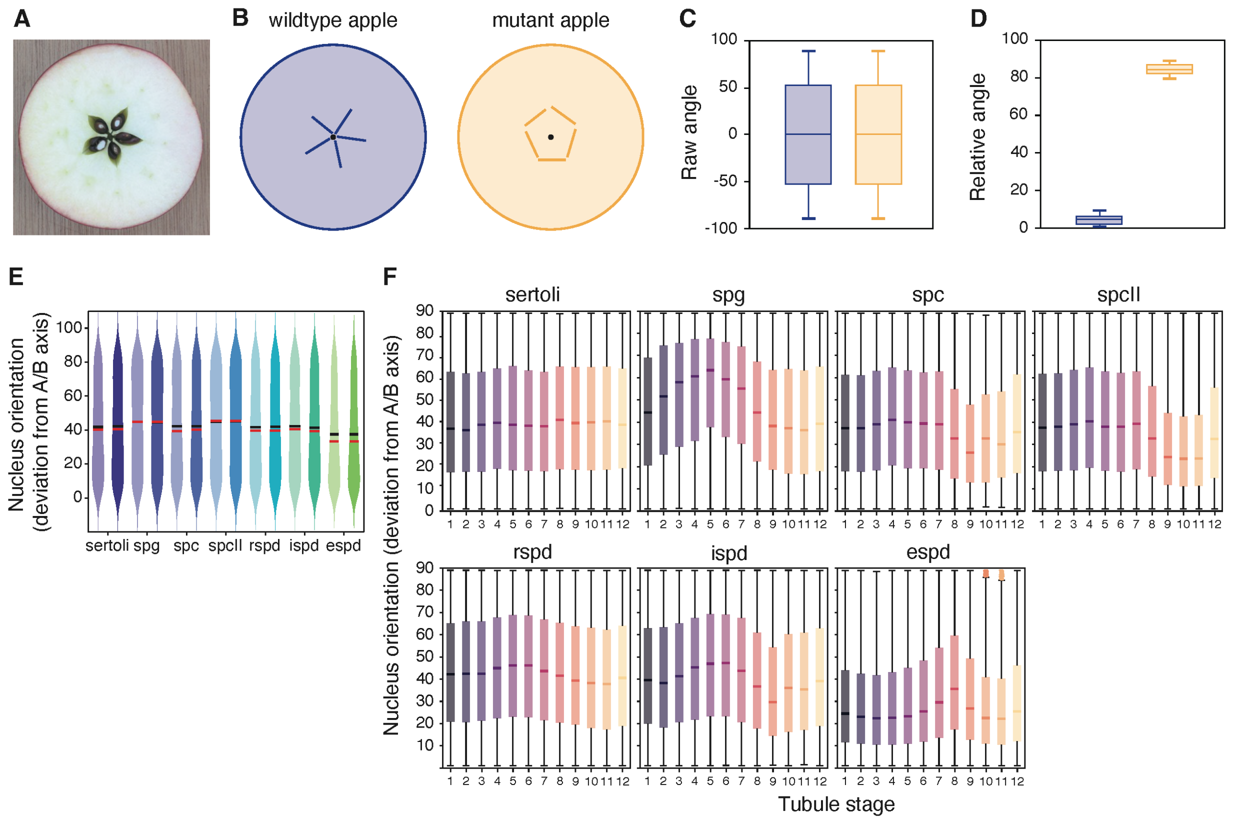


Fig. S3: Wildtype relative cell orientation plots. This statistic was developed to normalize cellular angle to a universal axis of reference for all tubules regardless of their cross-section. Cell angles are described as the offset to the radial vector, i.e. the vector between the tubule centroid and the cell centroid. The minimum possible value is 0°, corresponding to a cell whose major axis is parallel to the radial vector; the maximum possible value is 90°, indicating a cell whose major axis is orthogonal (circumferential). (A) We use an apple cross-section to illustrate the utility of this feature. (B) In the schematic for a wildtype apple (blue), the core compartments holding the seeds are oriented like a star. However, in a mutant apple (orange), suppose these vectors are now shifted so they are oriented in a circumferential direction. (C) Were we to measure the orientations of these vectors in (B) with no further context the distributions would be nearly identical. (D) However, if we were to measure their deviation from a radial vector, we can use this feature to quantify the effects of this mutation. (E-F) Actual data from wildtype tubules. (E) Orientation arranged by cell type. (F) Orientation arranged by both cell type and tubule stage.

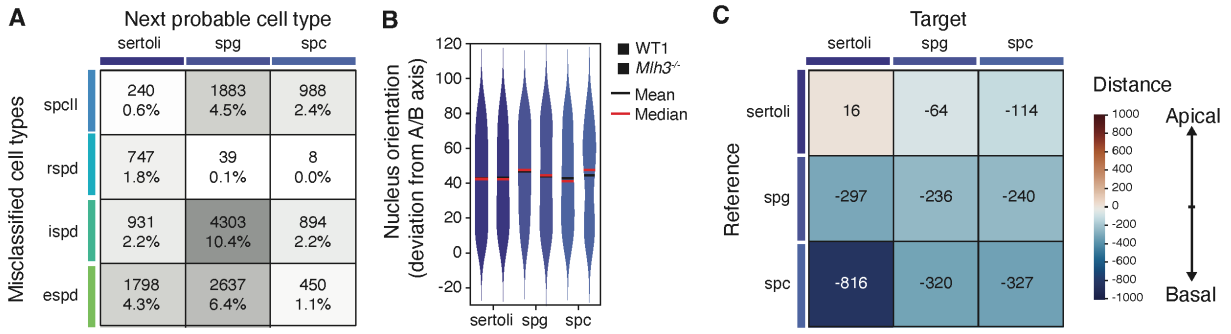


Fig. S4: Supplementary *Mlh3*$^{-/-}$ plots. (A) Classification errors in *Mlh3*$^{-/-}$ cells have no impact on overall result whether they are omitted or changed to next probable cell type. This assessment was used to ensure our results were not biased due to our cell assignment options. (B) Relative cell orientation did not significantly change for any *Mlh3*$^{-/-}$ cell type. (C) Nearest neighbor distances for the three cell classes in *Mlh3*$^{-/-}$. Most values are negative, indicating an increase in cell density along the basal membrane.
